# IFITM1 and IFITM3 cooperate to restrict virus entry in endolysosomes

**DOI:** 10.1101/2025.06.01.657267

**Authors:** Isaiah Wilt, Abigail A. Jolley, Kazi Rahman, Kin Kui Lai, Guoli Shi, Thorkell Andresson, Alex A. Compton

## Abstract

Interferon-induced transmembrane (IFITM) proteins are potent innate immune factors that restrict an array of viruses at the entry stage of infection. We previously characterized a GxxxG motif in the CD225 domain of human IFITM3 that mediates its multimerization, which is essential for the reduction of membrane fluidity by IFITM3 and for its antiviral activity against Influenza A virus. Here, using an unbiased approach coupling immunoprecipitation with mass spectrometry, we show that the GxxxG motif is also important for the interaction of IFITM3 with other proteins, including IFITM1. While IFITM1 is primarily regarded as a cell surface protein that restricts the entry of viruses fusing at the plasma membrane, this model is based mostly on overexpression studies and is at odds with some studies showing that it can restrict endocytic viruses. Here, we show that endogenous IFITM1 and IFITM3 co-reside in membranes of acidic late endosomes and lysosomes (endolysosomes) and form a protein-protein complex as determined by co-immunoprecipitation and proximity ligation assay. Knockdown of endogenous IFITM3 resulted in enhanced localization of IFITM1 at the plasma membrane, indicating that IFITM3 promotes IFITM1 localization to endolysosomes. To assess the antiviral protection conferred by endogenous IFITM1 and IFITM3 against viruses fusing at endolysosomal membranes, we measured cell entry mediated by the Influenza A fusogen hemagglutinin (HA). While knockdown of IFITM3 significantly boosted HA-mediated entry, combined knockdown of both IFITM3 and IFITM1 boosted entry even further. These results suggest that endogenous IFITM1 restricts Influenza A virus entry in a manner that is non-redundant with IFITM3, and that IFITM1 and IFITM3 inhibit virus entry in a cooperative manner.

## INTRODUCTION

Prior to the recruitment of innate and adaptive immune cells to sites of virus infection, the cell-intrinsic or cell-autonomous immune response is the first line of defense restricting infection within individual cells and inhibiting the spread of virus to neighboring bystander cells (1). Cell-intrinsic immunity is mediated by proteins encoded by interferon (IFN)-stimulated genes, many of which target different steps of the virus replication cycle (2). Several hundred IFN-stimulated genes are upregulated in response to type I, type II, and type III interferons, and only a fraction of these have been assigned functions or are understood on a mechanistic level (3). One group of IFN-stimulated genes that have been identified as broad-spectrum inhibitors of virus infection are the interferon-inducible transmembrane (IFITM) proteins consisting of IFITM1, IFITM2, and IFITM3 in humans. Primarily restricting the step of cellular entry, this group of restriction factors is believed to reduce membrane fusion between virus and cell by altering membrane fluidity and curvature (4, 5). Of the human IFITM proteins, IFITM3 exhibits potent antiviral activity against viruses that undergo membrane fusion in endosomes (6, 7), including Influenza A virus (8-11). Furthermore, the physiological importance of IFITM3 in vivo has been demonstrated by increased viral disease severity in Influenza-infected mice deficient for *Ifitm3* (12) and by increased disease severity in Influenza-infected humans carrying single nucleotide polymorphisms in *IFITM3* (13, 14). As a result, IFITM3 is the best-characterized member of the IFITM family, and its antiviral mechanism of action has been studied in greater detail.

IFITM1, IFITM2, and IFITM3 are paralogs that likely arose following multiple gene duplication events over evolutionary time, with IFITM3 serving as the most ancient, ancestral locus (15-17). Many studies have shown that IFITM proteins exhibit differences in subcellular localization. For example, some reports have suggested that IFITM1 is found primarily on the cell surface, while IFITM2 and IFITM3 are found primarily in early and late endosomes (10, 16, 18-24). It has been reported that viruses that enter cells by fusing at the plasma membrane, including respiratory syncytial virus, metapneumovirus, herpes simplex virus 1, and HIV-1 are restricted by IFITM1, and this has been attributed to its putative localization to the plasma membrane (25-27). However, other studies have reported that IFITM1 partially resides in intracellular membranes (26, 28-30) and that IFITM1 can inhibit virus entry taking place at sites other than the plasma membrane. Several studies have shown that IFITM1 exhibits antiviral activity against Influenza A virus, which is known to fuse with endosomal membranes in a pH-dependent manner (8, 31). Furthermore, IFITM1 restricts the filoviruses Ebola virus and Marburg virus, which enter cells by fusing with lysosomal membranes (23). Therefore, the apparent or assumed subcellular localization of IFITM1 in cells cannot fully explain virus sensitivity to IFITM1-mediated restriction. Moreover, most of work characterizing the properties of IFITM1 employ overexpression approaches in transformed human cell lines, often with an epitope tag, or even overexpression in non-human cells (28). The localization of endogenous IFITM1 protein that is constitutively expressed or interferon-upregulated in human cells, and how it relates to antiviral activity, has not been addressed. Moreover, it is unknown whether endogenous IFITM proteins interact with one another and whether these interactions in situ regulate their antiviral functions. While it was previously demonstrated that ectopic IFITM proteins could co-immunoprecipitate with one another (32), it was not addressed whether endogenous IFITM proteins interact and, if so, where these interactions occur on the subcellular level.

We previously found that IFITM3 forms homomultimers through a GxxxG motif found in its CD225 domain, and that this motif is essential for the antiviral functions of IFITM3 (33). Since the GxxxG motif is conserved in IFITM1 and IFITM2, it is likely that that IFITM1 and IFITM2 form homomultimers as well and that multimerization may be important for their respective functions. However, it remained a possibility that the GxxxG motif present in IFITM proteins enables interactions with other cellular proteins, including between different IFITM proteins. Here, we identified endogenous IFITM1 as a binding partner of IFITM3. Furthermore, we show that this interaction is dependent on glycine-95 motif of IFITM3, indicating that the conserved GxxxG motif is important for mediating both homo-and heteromultimerization among IFITM family members. By examining the interaction between endogenous IFITM3 and endogenous IFITM1 in diverse cell types, we found that these two proteins interact in acidic late endosomes and lysosomes (endolysosomes). Intriguingly, silencing of endogenous IFITM3 resulted in an accumulation of IFITM1 at the plasma membrane, revealing IFITM3 as an important regulator of IFITM1 localization. Therefore, the presence of IFITM3 may influence the antiviral properties of IFITM1 in the same cell. Since we found endogenous IFITM1 and IFITM3 to co-reside in endolysosomal membranes, we measured the extent to which silencing of IFITM3 alone or IFITM1 and IFITM3 together impacted cellular entry mediated by Influenza A hemagglutinin (HA). Our results demonstrated that both endogenous IFITM1 and IFITM3 act as barriers to the endocytic entry of Influenza A virus. These results provide insight into the cooperative roles played by IFITM proteins in cellular endomembranes. Moreover, they suggest that experiments involving the overexpression or silencing of individual IFITM genes should be interpreted cautiously.

## RESULTS

To identify interaction partners of IFITM3, including those dependent on the GxxxG motif of IFITM3 mediating homomultimerization and antiviral activity, we stably expressed FLAG-IFITM3 WT, FLAG-IFITM3 G95L, or Empty Vector in HEK293T cells, immunoprecipitated ectopic IFITM3 with anti-FLAG antibody from whole cell lysates, and identified co-immunoprecipitated proteins with proteomics (**Supplemental File 1**). Putative interactors of IFITM3 were qualified as those whose peptides were identified in the IFITM3 WT or IFITM3 G95L fractions or both but which were absent from the Empty Vector fraction. Furthermore, we sorted interactors that associated with IFITM3 WT to a greater extent than IFITM3 G95L to identify protein-protein interactions that depended on the presence of glycine-95. For each protein, overall abundance pulled down by IFITM3 WT was divided by abundance pulled down by IFITM3 G95L to derive an abundance ratio. Among the partners which preferentially pulled down with FLAG-IFITM3 WT was IFITM1 (**Figure 1A**). We found that IFITM3 WT pulled down approximately 9-fold more endogenous IFITM1 compared to IFITM3 G95L, indicating that G95 of IFITM3 promotes the interaction between IFITM3 and IFITM1 (**Figure 1A**).

**Figure 1.**
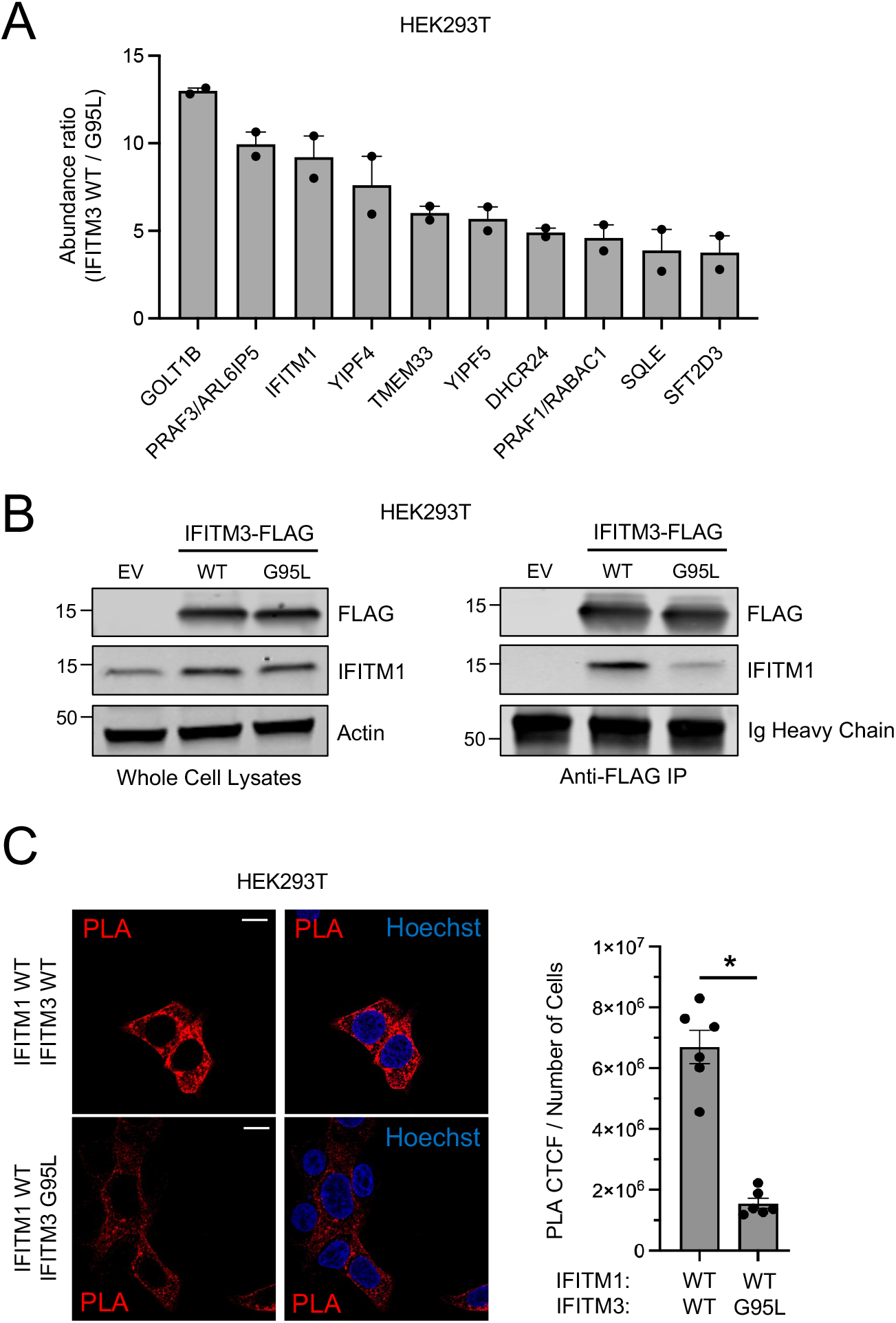
IFITM3 interacts with endogenous IFITM1 in a G95-dependent manner. (A) HEK293T cells stably expressing pQCXIP-FLAG-IFITM3 WT or pQCXIP-FLAG-IFITM3 G95L or PQCXIP-Empty Vector were lysed, and anti-FLAG immunoprecipitation was performed. Co-immunoprecipitated proteins were identified by mass spectrometry (MS). Abundance ratios were calculated whereby total abundance from immunoprecipitated FLAG-IFITM3 WT was divided by total abundance from FLAG-IFITM3 G95L. The mean abundance ratio plus standard error is shown for a subset of proteins that pulled down to a greater extent with IFITM3 WT relative to IFITM3 G95L. Symbols represent ratios obtained from duplicate MS runs. (B) Left: HEK293T cells stably expressing pQCXIP-FLAG-IFITM3 WT or pQCXIP-FLAG-IFITM3 G95L or pQCXIP-Empty Vector were lysed and whole cell lysates were subjected to SDS-PAGE and immunoblotting with anti-FLAG, anti-IFITM1, and anti-Actin (used as loading control). Right: FLAG-IFITM3 was immunoprecipitated with anti-FLAG, and IP fractions were subjected to SDS-PAGE and immunoblotting with anti-FLAG and anti-IFITM1 (immunoglobulin heavy chain served as loading control). Numbers and tick marks left of blots indicate position and size (in kilodaltons) of protein standard in ladder. (C) Left: HEK293T cells were transiently transfected with pQCXIP-FLAG-IFITM1 WT and pQCXIP-FLAG-IFITM3 WT or pQCXIP-FLAG-IFITM3 G95L, fixed and permeabilized, and proximity ligation assay was performed using anti-IFITM1 and anti-IFITM3 followed by confocal immunofluorescence microscopy. Nuclei were labeled with Hoechst. Scale bar = 15 microns. Right: The corrected total cell fluorescence was calculated and divided by number of cells and shown as mean plus standard error. Symbols represent fields of view containing 6-12 cells. Differences that were statistically significant between the indicated conditions as determined by student’s t test are indicated by (*) (*p* < 0.05). EV; empty vector. WT; wild type. Ig; immunoglobulin. PLA; proximity ligation assay. CTCF; corrected total cell fluorescence.

To verify that endogenous IFITM1 interacts with ectopic FLAG-IFITM3 in a G95-dependent manner, FLAG-IFITM3 WT and FLAG-IFITM3 G95L were immunoprecipitated with anti-FLAG and pull down of endogenous IFITM1 was determined by immunoblotting with anti-IFITM1 antibody. The specificity of anti-IFITM1 was determined by immunoblotting lysates from HeLa and HeLa IFITM3 knockout cells treated with type-I interferon or not (**Supplemental Figure 1A**). We found that endogenous IFITM1 pulled down with IFITM3 WT to a greater extent than with IFITM3 G95L, validating our proteomics result (**Figure 1B**). To determine whether IFITM3 and IFITM1 interact at the plasma membrane, intracellular membranes, or both, we performed proximity ligation assay (PLA) and confocal immunofluorescence microscopy using anti-IFITM3 and anti-IFITM1 in HEK293T transfected with IFITM1 and IFITM3. We found that PLA fluorescence was detected at or near the plasma membrane and was especially apparent in intracellular compartments in cells transfected with IFITM1 and IFITM3 WT. In contrast, fluorescence was significantly decreased in cells transfected with IFITM1 and IFITM3 G95L (**Figure 1C**). These results suggest that overexpressed IFITM3 interacts with both endogenous and overexpressed IFITM1 in a G95-dependent manner, and that this interaction can occur within endomembranes of cells.

While our findings reveal that IFITM1 and IFITM3 can interact with one another when one or both proteins are overexpressed in HEK293T cells, we next assessed the degree to which endogenously expressed IFITM1 and IFITM3 associate with each other when they are expressed constitutively or upregulated following interferon treatment. In HEK293T cells, the basal level of IFITM3 was undetectable while a limited quantity of IFITM1 was observed, while both IFITM1 and IFITM3 were significantly upregulated by type-I interferon (**Figure 2A**). When IFITM3 was immunoprecipitated from interferon-treated cells, we found that IFITM1 co-immunoprecipitated with it (**Figure 2A**). This interaction was validated in intact cells using proximity ligation assay, which demonstrated that endogenous IFITM1 and IFITM3 interact, in a direct or indirect manner, throughout the cytosolic space in interferon-treated cells (**Figure 2B**). To determine whether endogenous IFITM1 and IFITM3 associate in primary epithelial cells, we examined primary human nasal epithelia. Both IFITM1 and IFITM3 were detectable at a constitutive level in primary epithelia, and protein levels were enhanced following interferon treatment, and IFITM1 co-immunoprecipitated with IFITM3 in interferon-treated cells (**Figure 2C**). Therefore, our results suggest that endogenous IFITM1 and IFITM3 interact with one another in transformed and primary cells that are commonly used as targets for enveloped virus infection. We performed similar experiments in HeLa, since we and others previously demonstrated that IFITM proteins are constitutively expressed and restrict virus infections in these cells (34, 35). As expected, both IFITM1 and IFITM3 were observed in HeLa under basal conditions, with IFITM3 being more abundant (**Figure 2D**). Here, we observed that IFITM1 pulled down with immunoprecipitated IFITM3 in the absence and presence of interferon (**Figure 2D**). Confirming that IFITM1 and IFITM3 proteins interact constitutively in HeLa cells, abundant fluorescence was observed in perinuclear regions according to the proximity ligation assay, and this signal was abrogated in HeLa IFITM3 knockout cells (**Figure 2E**). Since IFITM2 is a close paralog of IFITM3, and since it is constitutively expressed in HeLa (albeit to a lesser extent than IFITM3) (**Supplemental Figure 1B**), we immunoprecipitated IFITM2 using a specific anti-IFITM2 antibody. We observed that both IFITM1 and IFITM3 co-immunoprecipitated with IFITM2 in HeLa cells, especially in interferon-treated cells (**Supplemental Figure 1B**).

**Figure 2.**
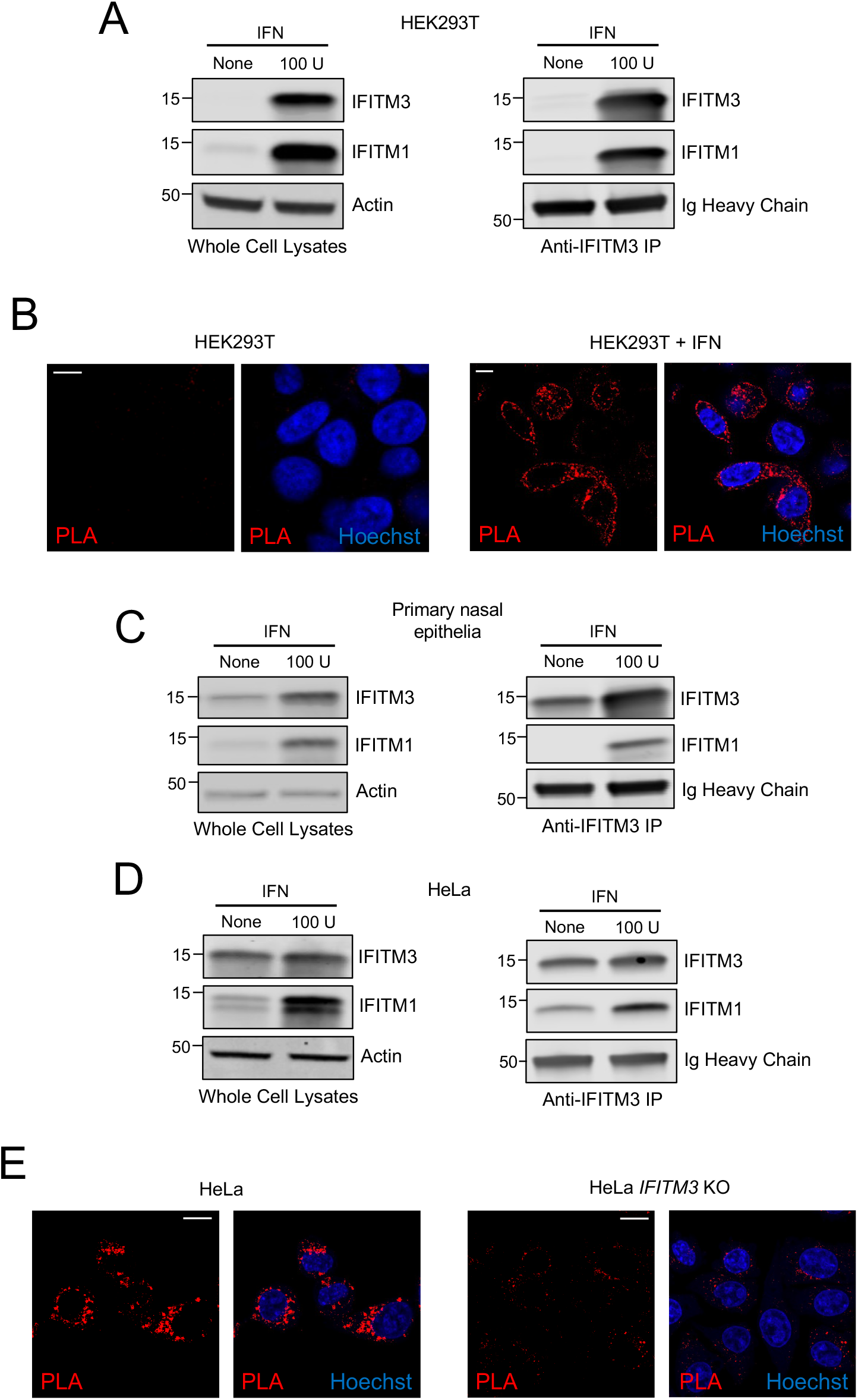
Endogenous IFITM3 and IFITM1 interact in multiple cell types. (A) Left: HEK293T cells were untreated or treated with 100 units type-I interferon (IFNb) for 18 hours and whole cell lysates were subjected to SDS-PAGE and immunoblotting with anti-IFITM3, anti-IFITM1, and anti-Actin (used as loading control). Right: IFITM3 was immunoprecipitated with anti-IFITM3 and IP fractions were subjected to SDS-PAGE and immunoblotting with anti-IFITM3 and anti-IFITM1 (immunoglobulin heavy chain served as loading control). (B) HEK293T cells were untreated or treated with 100 units IFNb for 18 hours, fixed and permeabilized, and proximity ligation assay was performed using anti-IFITM1 and anti-IFITM3 followed by confocal immunofluorescence microscopy. Nuclei were labeled with Hoechst. Scale bar = 10 microns. (C) Left: primary nasal epithelial cells were untreated or treated with 100 units IFNb for 18 hours and whole cell lysates were subjected to SDS-PAGE and immunoblotting with anti-IFITM3, anti-IFITM1, and anti-Actin (used as loading control). Right: IFITM3 was immunoprecipitated with anti-IFITM3 and IP fractions were subjected to SDS-PAGE and immunoblotting with anti-IFITM3 and anti-IFITM1 (immunoglobulin heavy chain was used as loading control). (D) Left: HeLa cells were untreated or treated with 100 units IFNb for 18 hours and whole cell lysates were subjected to SDS-PAGE and immunoblotting with anti-IFITM3, anti-IFITM1, and anti-Actin (used as loading control). Right: IFITM3 was immunoprecipitated with anti-IFITM3 and IP fractions were subjected to SDS-PAGE and immunoblotting with anti-IFITM3 and anti-IFITM1 (immunoglobulin heavy chain was used as loading control). Numbers and tick marks left of blots indicate position and size (in kilodaltons) of protein standard in ladder. (E) HeLa and HeLa IFITM3 knockout cells were fixed and permeabilized, and proximity ligation assay was performed using anti-IFITM1 and anti-IFITM3 followed by confocal immunofluorescence microscopy. Nuclei were labeled with Hoechst. Scale bar = 15 microns. IFN; interferon. U; units. KO; knockout.

We then performed confocal immunofluorescence microscopy to characterize the subcellular localization of endogenous IFITM1 and IFITM3. Partial colocalization between IFITM1 and IFITM3 at vesicular sites was observed in HeLa cells, and this was also the case in interferon-treated cells, although an apparent increase in IFITM1 at the cell periphery was observed (**Figure 3A**). In the same acquisitions, we assessed the position of IFITM1-positive vesicular compartments relative to Lysotracker, a marker of acidic late endosomes and lysosomes (endolysosomes). We found that a proportion of IFITM1 colocalized with Lysotracker in the absence and presence of interferon (**Figure 3B-C**). A portion of IFITM3 was also colocalized with Lysotracker, and the same Lysotracker-positive compartments contained both IFITM1 and IFITM3 (**Figure 3A**). Therefore, pools of IFITM1 and IFITM3 protein colocalize in endolysosomes in HeLa cells at constitutive and interferon induced levels. As a control to confirm the specificity of the anti-IFITM1 antibody, we knocked down endogenous IFITM3 with siRNA that selectively reduces IFITM3 protein. Surprisingly, while total IFITM1 levels were unaffected, there was an evident accumulation of IFITM1 at or near the cell periphery following knockdown of IFITM3 (**Figure 3A**). Furthermore, the colocalization between IFITM1 and Lysotracker was significantly reduced in IFITM3 knockdown cells (**Figure 3B-C**). These findings suggest that endogenous IFITM3 promotes the localization of endogenous IFITM1 to acidic endolysosomes.

**Figure 3.**
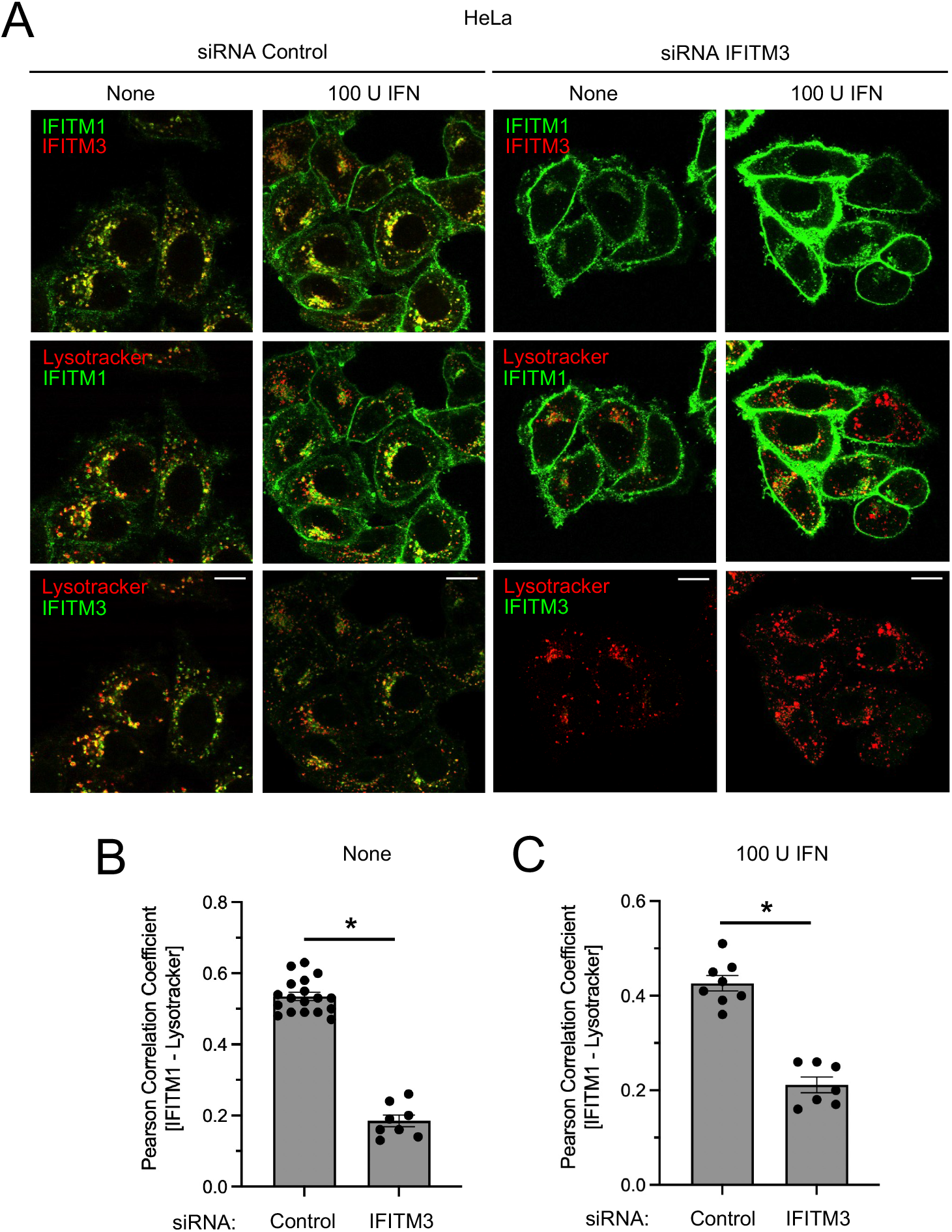
Endogenous IFITM1 and IFITM3 co-reside in endolysosomal membranes and the localization of IFITM1 to endolysosomes is IFITM3-dependent. (A) HeLa cells were transfected with control siRNA or siRNA targeting IFITM3 for 48 hours and subsequently treated with 100 units IFNb for 18 hours or left untreated. Cells were stained with Lysotracker, fixed and permeabilized, and immunostained with anti-IFITM1 and anti-IFITM3 followed by confocal immunofluorescence microscopy. Scale bar = 15 microns. (B) Colocalization between IFITM1 and Lysotracker in untreated cells was quantified by Pearson Correlation Coefficient and plotted as mean plus standard error. Symbols represent fields of view containing 5-12 cells. Differences that were statistically significant between the indicated conditions as determined by student’s t test are indicated by (*) (*p* < 0.05). (C) Colocalization between IFITM1 and Lysotracker in cells treated with 100 units IFNb was quantified by Pearson Correlation Coefficient and plotted as mean plus standard error. Symbols represent fields of view containing 5-12 cells. Differences that were statistically significant between the indicated conditions as determined by student’s t test are indicated by (*) (*p* < 0.05).

Having found that endogenous IFITM1 and IFITM3 interact and colocalize in endolysosomes, we sought to measure the capacity for both proteins to restrict an enveloped virus well-characterized to enter/fuse in endolysosomes. To this end, we produced vesicular stomatitis virus (VSV) pseudotyped with hemagglutinin (HA), the pH-dependent viral fusogen of Influenza A virus. The amount of VSV-HA infection of HeLa cells treated with control siRNA was normalized to 100. Following specific knockdown of IFITM3, VSV-HA infection was boosted approximately 4.5-fold (**Figure 4A-B**). However, when IFITM1 and IFITM3 were knocked down in combination, VSV-HA infection was boosted further (approximately 7-fold). We were unable to address the role of endogenous IFITM1 by itself because the siRNA targeting IFITM1 reduced not only IFITM1 protein but also IFITM3 protein (**Figure 4B**). Nonetheless, these results indicate that combined silencing of IFITM1 and IFITM3 result in increased cellular susceptibility to VSV-HA infection compared to silencing of IFITM3 alone. Therefore, endogenous IFITM1, which colocalizes with endogenous IFITM3 in endolysosomes, contributes to restriction of HA-mediated entry. Together with our observations that IFITM3 controls the localization of IFITM1 to endolysosomes, we propose that IFITM1 and IFITM3 cooperate to restrict virus entry in this compartment.

**Figure 4.**
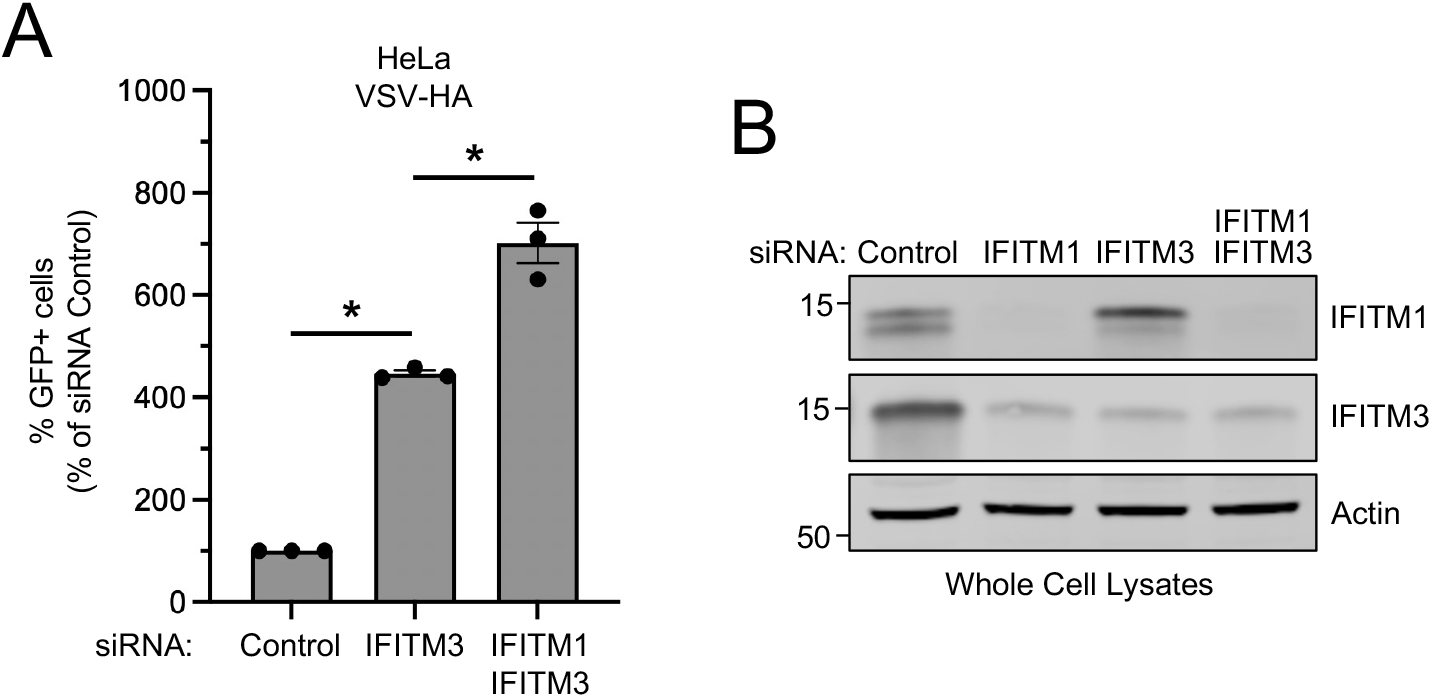
Endogenous IFITM1 and IFITM3 restrict HA-mediated virus entry. (A) HeLa cells were transfected with control siRNA, siRNA targeting IFITM3, or siRNA targeting IFITM3 plus siRNA targeting IFITM1 for 72 hours and inoculated with replication-incompetent VSV-HA pseudovirus. Infection was measured by GFP expression at 18 hours post-inoculation and plotted as mean plus standard error. Symbols represent three independent experiments (biological replicates). Differences that were statistically significant between the indicated conditions as determined by one-way ANOVA are indicated by (*) (*p* < 0.05). (B) HeLa cells were transfected with control siRNA, siRNA targeting IFITM1, siRNA targeting IFITM3, or siRNA targeting IFITM1 plus siRNA targeting IFITM3 for 72 hours. Cells were lysed and whole cell lysates were subjected to SDS-PAGE and immunoblotting with anti-IFITM1, anti-IFITM3, and anti-Actin (used as loading control). Numbers and tick marks left of blots indicate position and size (in kilodaltons) of protein standard in ladder. GFP; green fluorescent protein. VSV; vesicular stomatitis virus. HA; hemagglutinin.

## DISCUSSION

In this report, we show that endogenous IFITM1 associates with endogenous IFITM3 in multiple cell types, including primary human epithelial cells of the respiratory tract targeted by Influenza A Virus. The interaction between IFITM3 and IFITM1 seems to be consequential for IFITM1 function, since silencing of IFITM3 results in accumulation of IFITM1 at the plasma membrane. The observation that manipulation of IFITM3 levels can impact IFITM1 has important implications for the interpretation of past and future experiments that rely upon the overexpression or silencing of individual IFITM genes. Phenotypes generated by siRNA-mediated knockdown or CRISPR-Cas9-mediated knockout of IFITM3 may, in whole or in part, result from the indirect effect those manipulations have on IFITM1 protein. For example, the increase in Influenza A infection following IFITM3 knockdown may, at least in part, be due to the shift in IFITM1 localization to the plasma membrane. This is a distinct possibility, since IFITM1 overexpression has been reported to restrict Influenza A virus entry, and since we show here that both endogenous IFITM1 and endogenous IFITM3 contribute to restriction of Influenza A virus entry, as indicated by the combined effect of IFITM1 and IFITM3 knockdown. However, these results likely underestimate the antiviral role of endogenous IFITM1, since we also show that loss of endogenous IFITM3 results in relocalization of IFITM1 to the cell surface. One study showed that combined knockout of IFITM1, IFITM2, and IFITM3 in HeLa did not result in more elevated Influenza A infection compared to IFITM2 and IFITM3 knockout alone (36), and this was interpreted as suggesting that IFITM1 does not have an antiviral role to play against Influenza A virus infection in HeLa cells. However, the lack of additional effect resulting from IFITM1 might be mitigated by the fact that IFITM1 is mislocalized in the IFITM2/3 knockout cells. Also, it is unknown how IFITM2, or the lack thereof, may influence the antiviral activity of IFITM1.

Our results imply that the localization and activities of IFITM1 are influenced by the relative abundance of IFITM3 in the same cell. Since the constitutive level of IFITM3 protein present in different cell types may vary, the localization and activity of IFITM1 may also vary between cell types as a result. Indeed, it has been shown that IFITM1 exhibits cell-type dependent effects during certain virus infections (6, 7, 28, 37). Furthermore, it is likely that interferons can impact IFITM1 localization and function not only by upregulating IFITM1 itself, but by upregulation of IFITM3. In addition to the impact of IFITM3 on IFITM1 localization and function, it is also possible that IFITM1 contributes to phenotypes resulting from IFITM3 overexpression, since we show that overexpressed IFITM3 interacts with endogenous IFITM1 (**Figure 1B**). These findings provide a new, expanded framework for interpreting the effects of single IFITM knockdown or single IFITM overexpression.

What remains to be determined is the mechanism by which IFITM3 promotes the endolysosomal localization of IFITM1. It seems clear that IFITM3 does not promote the targeted degradation of IFITM1 in endolysosomes, since the presence of IFITM3 was not accompanied by decreased total IFITM1 protein levels (**Figure 1B and Figure 4B**). It will be important to ascertain how mutations that disrupt IFITM3 localization or function, including sites which are post-translationally modified, impact binding to IFITM1 and the localization of IFITM1 to endolysosomes. Furthermore, since it was previously shown that IFITM1 contains an endocytic sorting signal in its carboxy terminus which enables AP-3-dependent targeting to endoylysosomes (26, 29, 30), it is worth examining whether IFITM3 regulates the exposure or recognition of this sorting signal. It is possible that, in the presence of IFITM3, the carboxy terminal sorting signal of IFITM1 is stabilized or otherwise made more accessible to AP-3. Interestingly, we found that our IFITM1-specific antibody recognized two populations of endogenous IFITM1 in HeLa cells. Upon knockdown of IFITM3, the more slowly migrating population of IFITM1, which may correspond to a longer variant of polypeptide, predominated (**Figure 4B**). This may signify that IFITM3, by enabling the localization of IFITM1 to endolysosomes, facilitates the cleavage of the carboxy terminus of IFITM1 (which is expected to face the lysosomal lumen (19, 30)) by lysosomal hydrolases. IFITM3 may directly promote IFITM1 localization to endolysosomes through a protein-protein interaction. However, it is also possible that IFITM3 promotes the delivery of IFITM1 to endolysosomes through a more indirect mechanism—we recently demonstrated that IFITM3 promotes the lysosomal delivery of endocytic cargo by regulating late endosome fusion with lysosomes (38).

IFITM2 is the product of a recent gene duplication of IFITM3 in the ancestor of Homininae (humans, chimpanzees, and gorillas) (16). Since we show that endogenous IFITM2 co-immunoprecipitates with both IFITM1 and IFITM3, it may provide an additional regulatory layer to the function of IFITM1. Future work will focus on determining whether IFITM2 and IFITM3 perform redundant or distinct roles with regards to regulating the subcellular localization of IFITM1. Overall, our results reveal how regulatory relationships arose between closely related antiviral effectors that originated through multiple gene duplication events. Since unique duplications of IFITM genes have occurred in many species outside of humans (16, 39), resulting in unique repertoires of IFITM proteins that vary by species, there likely exist diverse mechanisms of cooperativity and regulation between IFITM proteins that remain to be discovered.

## Supporting information

IP-MS Abundance Results

## FIGURE LEGENDS

## Supplemental Figure 1

(A) HeLa and HeLa IFITM3 knockout cells were untreated or treated with 100 units IFNb for 18 hours and lysed. Whole cell lysates were subjected to SDS-PAGE and immunoblotting with anti-IFITM3, anti-IFITM1, and anti-Actin (used as loading control). (B) Left: HeLa cells were untreated or treated with 100 units or 500 units IFNb for 18 hours and lysed. Whole cell lysates were subjected to SDS-PAGE and immunoblotting with anti-IFITM1, anti-IFITM3, anti-IFITM2, and anti-Actin (used as loading control). Right: IFITM2 was immunoprecipitated with anti-IFITM2 and IP fractions were subjected to SDS-PAGE and immunoblotting with anti-IFITM1, anti-IFITM3, and anti-IFITM2 (immunoglobulin heavy chain was used as loading control). Numbers and tick marks left of blots indicate position and size (in kilodaltons) of protein standard in ladder.

## METHODS

### Cells and cell culture reagents

HEK293T (ATCC; CRL-3216), and HeLa (ATCC; CCL-2) were cultured in Dulbecco’s modified Eagle’s medium (DMEM, Gibco) supplemented with 10% heat-inactivated fetal bovine serum (FBS, Hyclone) and 1% penicillin-streptomycin (Gibco) at 37°C with 5% CO_2_. HeLa IFITM3 KO cells were previously described (38). HEK293T stably expressing pQCXIP-Empty Vector, pQCXIP-FLAG-IFITM3 WT, and pQCXIP-FLAG-IFITM3 G95L were previously described (33). Human primary nasal epithelial cells (Epithelix; EP51AB) were cultured in hAEC Culture Medium (Epithelix; EP09AM) at 37°C with 5% CO_2_. Human interferon beta 1a was obtained from PBL Assay Science (11415-1). Lysotracker Deep Red was obtained from Thermo Fisher (L12492).

### SDS-PAG, Western blot analysis, and immunoprecipitation

Whole cell lysis was performed using a buffer consisting of 20 mM HEPES, 150 mM NaCl, 1 mM EDTA, and 1% Triton X-100 (Sigma; X100) containing Halt protease inhibitor cocktail, EDTA-free (Thermo Fisher; 78425). Lysis was performed on ice for 30 mins prior to centrifugation at 12,000 rpm for 10 mins at 4°C, and supernatants were harvested. Protein quantity was measured using the Protein Assay (Bio-Rad; 5000001). Lysates were mixed with NuPAGE Reducing Agent (Invitrogen; NP0009) and Protein Loading Buffer (Li-COR; 928-40004) and loaded into Criterion XT 12% polyacrylamide Bis-Tris gels (Bio-Rad; 3450117). SDS-PAGE was performed with NuPAGE MES SDS Running Buffer (Invitrogen; NP0002). Proteins were transferred to Amersham Protein Nitrocellulose Membrane, pore size 0.2 microns (GE Healthcare; 10600004).

The following primary antibodies were used: anti-IFITM3 antibody (Abcam; ab109429), anti-FLAG antibody (Sigma; F7425), anti-IFITM1 (Proteintech; 60074-1-Ig), anti-IFITM2 (Proteintech; 66137-1-Ig), anti-actin (Santa Cruz Biotechnology; sc-47778). The following secondary antibodies were used: goat anti-mouse IRDye 800CW (Li-COR; 926-32210), goat anti-mouse IRDye 680RD (Li-COR; 926-68070), goat anti-rabbit 800CW (Li-COR; 926-32211), goat anti-rabbit 680RD (Li-COR; 926-68071). For immunoprecipitation, anti-IFITM3 antibody (Abcam; ab109429) or anti-FLAG antibody (Sigma; F7425) was added to 300 *μ*g whole cell lysates, and the protein-antibody mixture was incubated for 3 hours at 4°C. 15 *μ*L of Dynabeads Protein G (Invitrogen; 10007D) (pre-washed in cell lysis buffer) was added to the protein-antibody mixture and incubated by rotating for 1 hour at 4°C, and reaction mixtures were centrifuged at 1000 x g for 3 mins at 4°C. The supernatant was removed, and the pelleted beads were washed three times with cell lysis buffer. The bead fraction was resuspended with NuPAGE Reducing Agent (Invitrogen; NP0009) and Protein Loading Buffer (Li-COR; 928-40004), boiled for 5 mins at 95°C, and loaded into Criterion XT 12% polyacrylamide Bis-Tris gels (Bio-Rad; 3450117). Images were obtained with the Li-COR Odyssey CLx and analysis was performed with ImageStudio Lite software (Li-COR). PageRuler Prestained Protein Ladder, 10-180 kDa was used as protein standard (Thermo Fisher; 26616).

### Confocal immunofluorescence microscopy

Cells were seeded in 8-well Mu chamber slides (Ibidi; 80826) at 15,000 cells per well. For HEK293T cells, chamber slides were coated with 3 mg/mL PureCol for one hour at room temperature and allowed to dry prior to cell seeding. Living cells were stained with Lysotracker Deep Red (Thermo Fisher; L12492) at a final concentration of 50 nM in complete DMEM for 30 mins at 37°C. Cells were washed with D-PBS (Gibco) and fixed/permeabilized with Cytofix/Cytoperm Solution (BD; 554722) for 10 mins at room temperature. Cells were washed with Cyto Perm/Wash Buffer (BD; 554723) and blocked in Intercept Blocking Buffer (PBS) (Licor; 927-70001) for one hour at room temperature. Primary antibodies (rabbit anti-IFITM3 (Abcam; ab109429) and mouse anti-IFITM1 (Proteintech; 60074-1-Ig) were diluted 200-fold in Intercept T20 Antibody Diluent (PBS) (Licor; 927-75001) and cells were incubated for 3 hours at room temperature. Cells were washed twice with Cyto Perm/Wash Buffer. Secondary antibodies (goat anti-mouse IgG (H+L) Secondary Antibody, Alexa Fluor 488 (Invitrogen; A11001) and goat anti-rabbit IgG (H+L) Secondary Antibody, Alexa Fluor 555 (Invitrogen; A21428)) were diluted 300-fold in Intercept T20 Antibody Diluent and cells were incubated for one hour at room temperature. Cells were washed twice with Cyto Perm/Wash Buffer and stained with Hoechst 33342 (Thermo Fisher; H3570) diluted 3000-fold in D-PBS for 5 mins. Cells were washed with Cyto Perm/Wash Buffer and imaged in D-PBS. Image acquisition was performed with a Leica Stellaris confocal fluorescence microscope. Images were analyzed and processed using Fiji (Image J).

### Immunoprecipitation-mass spectrometry (IP-MS)

HEK293T stably expressing pQCXIP-FLAG-IFITM3 WT, pQCXIP-FLAG-IFITM3 G95L, or pQCXIP-Empty Vector were lysed (33), and immunoprecipitation of FLAG-IFITM3 was performed with anti-FLAG (M2) (Sigma; F1804) according to the protocol indicated above. Protein Digestion and TMT labeling. Immunoprecipitated fractions were lysed in 50 mM HEPES, pH 8.0 and 8M urea followed by sonication. Lysates were clarified by centrifugation and protein concentration was quantified using BCA protein estimation kit (Thermo Fisher). One hundred micrograms of lysate were alkylated and digested by addition of trypsin at a ratio of 1:50 (Promega) and incubating overnight at 37^°^C. Digestion was acidified by adding formic acid (FA) to a final concentration of 1% and desalted using peptide desalting columns (Thermo Fisher) according to manufacturer’s protocol. Peptides were eluted from the columns using 50% ACN/0.1% FA, dried in a speedvac, and kept frozen at -20^°^C until further analysis. For TMT labeling, 15 ug of each sample was reconstituted in 50 uL of 50 mM HEPES, pH 8.0, and 75 ug of TMTpro label (Thermo Fisher) in 100% ACN was added to each sample. After incubating the mixture for 1 hr at room temperature with occasional mixing, the reaction was terminated by adding 8 uL of 5% hydroxylamine. The peptide samples for each condition were pooled and cleaned using peptide desalting columns (Thermo Fisher). High pH reverse phase fractionation. The first dimensional separation of the peptides was performed using a Waters Acquity UPLC system coupled with a fluorescence detector (Waters) using a 150 mm x 3.0 mm Xbridge Peptide BEM^™^ 2.5 um C18 column (Waters) operating at 0.35 mL/min. The dried peptides were reconstituted in 100 uL of mobile phase A solvent (3 mM ammonium bicarbonate, pH 8.0). Mobile phase B was 100% acetonitrile (Thermo Fisher). The column was washed with mobile phase A for 10 min followed by gradient elution 0-50% B (10-60 min) and 50-75 %B (60-70 min). The fractions were collected every minute. These 60 fractions were pooled into 24 fractions. The fractions were vacuum centrifuged to dryness and stored at -80^°^C until analysis by mass spectrometry. Mass Spectrometry acquisition and data analysis. The dried peptide fractions were reconstituted in 0.1% TFA and subjected to nanoflow liquid chromatography (Thermo Ultimate^™^ 3000RSLC nano LC system, Thermo Scientific) coupled to an Orbitrap Eclipse mass spectrometer (Thermo Scientific). Peptides were separated using a low pH gradient using 5-50% ACN over 120 minutes in mobile phase containing 0.1% formic acid at 300 nL/min flow rate. MS scans were performed in the Orbitrap analyser at a resolution of 120,000 with an ion accumulation target set at 4e^5^ and max IT set at 50ms over a mass range of 400-1600 m/z. Ions with determined charge states between 2 and 5 were selected for MS2 scans in the ion trap with CID fragmentation (Turbo; NCE 35%; maximum injection time 35□ms; AGC 1□×□10^4^). The spectra were searched using the Real Time Search Node in the tune file using human Uniprot database using Comet search algorithm with TMT16 plex (304.2071Da) set as a static modification of lysine and the N-termini of the peptide. Carbamidomethylation of cysteine residues (+57.0214□Da) was set as a static modification, while oxidation of methionine residues (+15.9949□Da) was set up as dynamic modification. For the selected peptide, an SPS–MS3 scan was performed using up to 10 *b*- and *y*-type fragment ions as precursors in an Orbitrap at 50,000 resolution with a normalized AGC set at 500 followed by maximum injection time set as “Auto” with a normalized collision energy setting of 65. Acquired MS/MS spectra were searched against a human Uniprot protein database along with a contaminant protein database, using a SEQUEST and percolator validator algorithms in the Proteome Discoverer 2.4 software (Thermo Scientific). The precursor ion tolerance was set at 10 ppm and the fragment ions tolerance was set at 0.02 Da along with methionine oxidation included as dynamic modification. Carbamidomethylation of cysteine residues and TMT16 plex (304.2071Da) was set as a static modification of lysine and the N-termini of the peptide. Trypsin was specified as the proteolytic enzyme, with up to 2 missed cleavage sites allowed. Searches used a reverse sequence decoy strategy to control for the false peptide discovery and identifications were validated using percolator software. Reporter ion intensities were adjusted to correct for the impurities according to the manufacturer’s specification and the abundances of the proteins were quantified using the summation of the reporter ions for all identified peptides. The reporter abundances were normalized across all the channels to account for equal peptide loading.

### Proximity ligation assay

In situ proximity ligation assay (PLA) was performed with the Duolink In Situ Red Starter Kit Mouse/Rabbit (Sigma; DUO92101) according to the manufacturer’s protocol. Cells were seeded in 8-well Mu chamber slides (Ibidi; 80826) at 15,000 cells per well. For HEK293T cells, chamber slides were coated with 3 mg/mL PureCol for one hour at room temperature and allowed to dry prior to cell seeding. Cells were fixed/permeabilized with Cytofix/Cytoperm Solution (BD; B554714) for 5 min and blocked with Duolink Blocking Solution (1X) for 1 hour at 37°C. Cells were then incubated with primary antibodies (rabbit anti-IFITM3 (Abcam; ab109429) and mouse anti-IFITM1 (Proteintech; 60074-1-Ig)) for 1 hour at room temperature. Cells were washed twice with Buffer A and subsequently incubated with the probes affinity purified Donkey anti-Rabbit IgG (anti-Rabbit PLUS, Sigma; DUO92002) and affinity purified Donkey anti-Mouse IgG (anti-Mouse PLUS, Sigma; DUO92004) for 1 hour at 37°C). After washing cells twice with Buffer A, oligonucleotide ligation was performed for 30 min at 37°C. Cells were washed two additional times with Buffer A, followed by incubation with amplification stock solution for 100 min at 37°C. After washing twice with Buffer B, Hoechst 33342 (Thermo Fisher; H3570) was added for 5 min at room temperature to label nuclei. Image acquisition was performed with a Leica Stellaris confocal fluorescence microscope.

### Pseudovirus production and infection

Viral RNA was extracted from HEK293T cells infected with Influenza A/PR/8/34 (PR8; H1N1) using the RNeasy Mini Kit (Qiagen). Two-step RT-PCR was performed to specifically amplify viral hemagglutinin (HA). Briefly, 1 μg or viral RNA was reverse transcribed into cDNA using Transcriptor First Strand cDNA Synthesis Kit (Roche) using the Uni12 primer (40).

Subsequently, HA cDNA was amplified using Platinum SuperFi II DNA Polymerase (Thermo Fisher) with the following primers: 5’-aaaaaaaggtaccgaaatgaaggcaaacctactaggtcctgttat-3’ and 5’-aaaaaaagcggccgctcagatgcatattctgcactgcaaa-3’. HA was cloned into pcDNA3.1(+) using KpnI and NotI restriction sites to produce pcDNA3.1-HA. HEK293T cells were seeded in a 10 cm dish and transfected with 8 μg pcDNA3.1-HA and 8 μg pcDNA3.1-NA (encoding neuraminidase, previously described (41)) using TransIT-293 Transfection Reagent (Mirus) and incubated at 37°C. Forty-eight hours post-transfection, cell culture medium was removed and 1 mL VSV-luc/GFP “seed particles” were added (42). Two hours later, cell culture medium was removed, and cells were washed three times with serum-free DMEM. Complete DMEM was added, and cells were incubated at 37°C for an additional 16 hours. Pseudovirus-containing supernatants were harvested, clarified by centrifugation at 500 x g for 5 min, filtered through a 0.22 micron filter. HeLa cells were incubated with pseudovirus for 18 hours at 37°C and GFP+ cells were enumerated with a Tecan Spark plate reader.

### siRNA transfection

For immunofluorescence experiments, HeLa cells were transfected with 20 nM Silencer Select siRNA (Thermo Fisher) (Negative Control #1 (4390844) or 20 nM IFITM3 (4392421; s195035)) using Lipofectamine RNAiMAX according to manufacturer’s instructions. siRNA and RNAiMAX were diluted in OptiMEM Reduced Serum Medium (Gibco). Cells were cultured at 37°C for 72 hours post-transfection followed by fixation, permeabilization, and immunostaining for confocal immunofluorescence microscopy. For VSV-HA pseudovirus infection experiments, HeLa cells were transfected with 40 nM Silencer Select siRNA (Thermo Fisher) (Negative Control #1 (4390844), 20 nM IFITM1 (4392421; s16192), 20 nM IFITM3 (4392421; s195035)) or 20 nM IFITM1 (4392421; s16192) plus 20 nM IFITM3 (4392421; s195035) using Lipofectamine RNAiMAX according to manufacturer’s instructions. siRNA and RNAiMAX were diluted in OptiMEM Reduced Serum Medium (Gibco). Cells were cultured at 37°C for 72 hours post-transfection followed by addition of VSV-HA pseudovirus.

### Statistical analysis

Tests for statistical significance were performed in GraphPad Prism.

## FUNDING

This work was funded by the Intramural Research Program, Center for Cancer Research, National Cancer Institute, National Institutes of Health.

## ACKNOWLEDGEMENTS

We would like to thank Mahesh Agarwal for assistance with flow cytometry and Bhabadeb Chowdhury for assistance with plasmid DNA amplification and purification.

## Notes

### Competing Interest Statement

The authors have declared no competing interest.

