## Supplementary figures and images for "IFITM1 and IFITM3 cooperate to restrict virus entry in endolysosomes"

### Supplemental Figure 1

Supplemental Figure 1

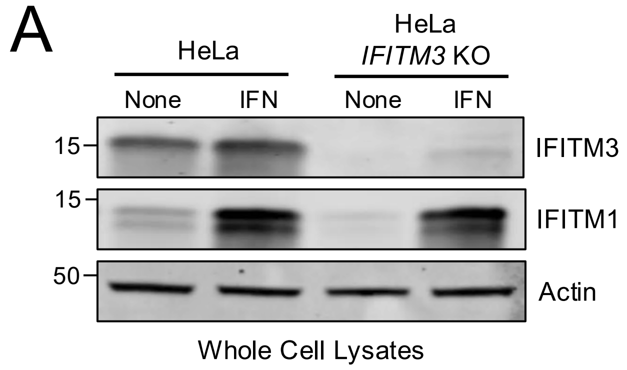


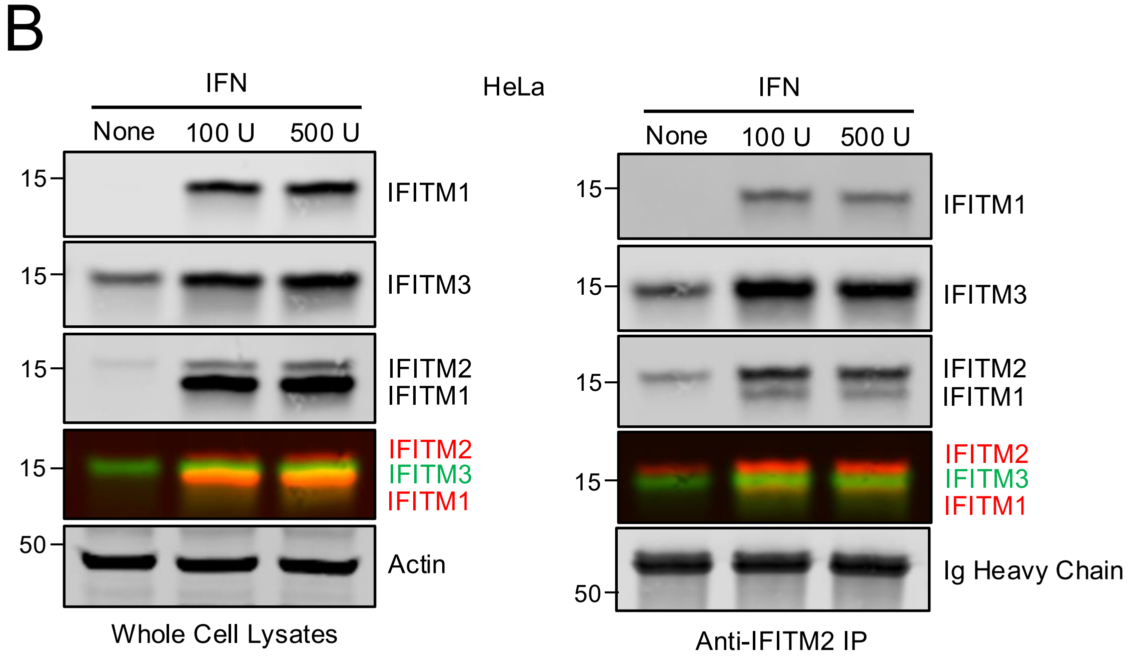
